# Spacer sequences separating transcription factor binding motifs set enhancer quality and strength

**DOI:** 10.1101/098830

**Authors:** Marion Guéroult-Bellone, Kazuhiro R. Nitta, Willi Kari, Edwin Jacox, Rémy Beulé Dauzat, Renaud Vincentelli, Carine Diarra, Ute Rothbächer, Christelle Dantec, Christian Cambillau, Jacques Piette, Patrick Lemaire

**Affiliations:** Centre de Recherche en Biologie cellulaire de Montpellier, UMR 5237, CNRS-Université de Montpellier, 1919 route de Mende, 34293 Montpellier, France; Institut de Biologie du Dévelopment de Marseille- IBDM, UMR7288 CNRS - Aix-Marseille Univ. - Case 907, 163 Avenue de Luminy 13288 Marseille CEDEX 09 FRANCE; Current address: Department of Evolution and Developmental Biology, Zoological Institute, University Innsbruck, Technikerstrasse 25, A-6020 Innsbruck, Austria.; Architecture et Fonction des Macromolécules Biologiques-AFMB - UMR7257 CNRS - Aix-Marseille Univ. - Case 932, 163 Avenue de Luminy 13288 Marseille CEDEX 09 FRANCE

**Keywords:** *Ciona intestinalis*, gene regulation, development, enhancer, transcription factor, DNA shape, Ets, Gata, Otx, ascidian, transcription factor affinity

## Abstract

Only a minority of the many genomic clusters of transcription factor binding motifs (TFBM) act as transcriptional enhancers. To identify determinants of enhancer activity, we randomized the spacer sequences separating the ETS and GATA sites of the early neural enhancer of the tunicate *Ciona intestinalis Otx* gene. We show that spacer sequence randomization affects the level of activity of the enhancer, in part through distal effects on the affinity of the transcription factors for their binding sites. A possible mechanism is suggested by the observation that the shape of the DNA helix within the TFBM can be affected by mutation of flanking bases that modulate transcription factor affinity. Strikingly, dormant genomic clusters of ETS and GATA sites are awakened by most instances of spacer randomization, suggesting that the sequence of naturally-occurring spacers ensures the dormancy of a majority of the large reservoir of TFBM clusters present in a metazoan genome.

## INTRODUCTION

Enhancers play a fundamental role in development, homeostasis, evolution and disease (1, 2). They act as scaffolding platforms for transcription factors and are generally composed of clusters of several binding sites for at least two transcription factors (3). The degree of constraints on the spacing, order and orientation of transcription factor binding sites is variable, with a majority of enhancers active during animal development showing little constraints (4). In spite of this apparent flexibility, we do not understand the determinants of enhancer activity and it remains very difficult to rationally engineer synthetic enhancers from the sole knowledge of upstream transcription factor binding sites (5).

The a-element of the ascidian *Ciona intestinalis* is one of the best-characterized chordate enhancers (6–9). This short (55 bp) enhancer drives the embryonic expression of the *Otx* gene from the late 32-cell stage in two animal neural lineages, a6.5 and b6.5 (Figure 1A), in response to the FGF9/16/20 neural inducer (6). This element is also weakly active in the posterior muscle lineage (B6.4) and in the neural progeny of the a6.7 cell pair, two territories that also express *Otx* (Figure 1A).

**Figure 1:**
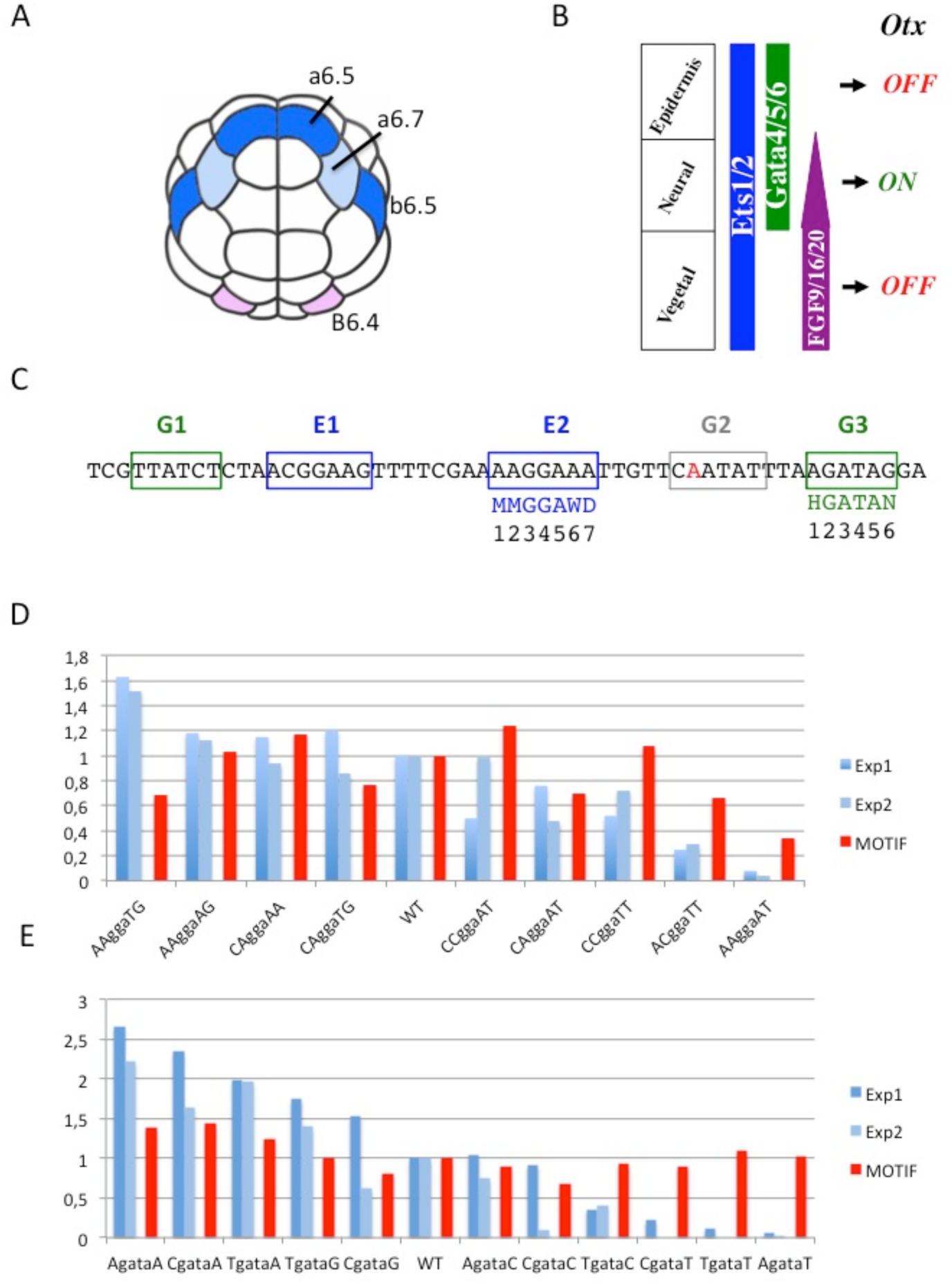
Influence of point mutations of the E2 and G3 sites on *in silico* transcription factor binding scores and *in vivo* enhancer activity. A) 32-cell stage embryo with cells in which the a-element is strongly (blue) or weakly active (light blue for neural and pink for muscle lineage). B) Neural induction of the a-element by the combined activity of Gata4/5/6 and Ets1/2. See text for details. Adapted from (6). C) a-element sequence showing the mutations in the ETS (blue) and GATA (green) sites tested *in vivo*. The inactivated (G>A in red) G2 site is in grey. M stands for A or C, W for T or A, H for A, T or C and N for any base. D) Effect of E2 mutations. Comparison of relative *in vivo* enhancer activity in two independent experiments (two shades of blue) and *in silico* predicted binding (red) of Ets1/2 for the E2 mutants. E) Effect of G3 mutations. Comparison of relative *in vivo* enhancer activity in two independent experiments (two shades of blue) and *in silico* predicted binding (red) of Gata4/5/6 for the G3 mutants. Activity is relative to the WT a-element.

The *cis*-regulatory logic driving the activity of this element in neural lineages has been characterized in detail (Figure 1B). Two maternal transcription factors, Ets1/2 and Gata4/5/6, cooperate to mediate FGF inducibility and tissue specificity, respectively (6,7). Binding of the ubiquitous Ets1/2 transcription factor to its sites drives expression in FGF-responding cells across all germ layers, while binding of the animal determinant Gata4/5/6 restricts this activation to the animal territories (Figure 1B). Mutational inactivation of individual ETS and GATA sites indicate that binding of Gata4/5/6 and Ets1/2 to two sites each is crucial for a- element activity (Figure 1C; 8).

The spacing and orientation of ETS and GATA binding sites does not seem to play a major role in a-element activity (8). In spite of this apparent flexibility, only a minority of the numerous *Ciona* genomic clusters containing at least 2 ETS and 2 GATA binding motifs have enhancer activity (8). A recent study of the a-element proposed that the major determinants of enhancer activity are included in the octamers composed of the core recognition tetramer for Ets1/2 (GGAA) and Gata4/5/6 (GATA) flanked by two adjacent nucleotides on either side (9). Dinucleotide repeat motifs required for enhancer function in addition to the transcription factor motifs were also characterized in *Drosophila* (10). In addition, transcription factor binding is, in part, determined by nucleotides lying outside of the DNA sequence directly contacted by the factor (11–14). Here, we combine *in vivo* and *in vitro* studies in a thorough analysis of the sequence determinants of the activity of the a-element enhancer and of other *Ciona* potential early neural enhancers responding to the same *cis*-regulatory logic.

## RESULTS

### Contribution of the bases contacted by Transcription Factors to enhancer activity

Our aim was to determine the respective roles of transcription factor binding sites and spacer sequences in a-element enhancer activity. For the sake of simplification, we used an a-element variant in which the G2 site was inactivated by point mutation, as this site was shown to be dispensable for enhancer activity (Figure 1C; 8). We first analysed the influence of the stretch of DNA directly contacted by Ets1/2 and Gata4/5/6 on the *in vivo* enhancer activity of the a-element. Published crystal structures for mammalian homologs bound to DNA (Supplementary Figure 1) suggest that Gata4/5/6 directly contacts 6 nucleotides: a central “GATA” core motif flanked on either side by one nucleotide (14, 15, 16). The contacts established by Ets1/2 span 7 nucleotides centered on GGA (17, 18, 19). To assess the relative importance of the nucleotides flanking the invariant “GGA” core of E2 and the “GATA” core of G3 (Figure 1C) we compared the activity of several combinations of point mutations by scoring LacZ staining in the a6.5 and b6.5 lineages in 112-cell stage embryos electroporated with reporter constructs (Figure 1D, E; see material and methods).

As expected, changes in the sequences of E2 and G3 quantitatively affected activity levels of the a-element in a6.5 and/or b6.5 lineages, while qualitatively preserving its spatial pattern of activity (not shown). In response to alteration of either binding site, the output levels of variant enhancers ranged from an almost complete inactivity to stronger than WT levels (Figure 1D, E; blue bars).

Surprisingly, k-mer based *in silico*-predicted affinities of the E2 and G3 variant octamer sites, derived from Selex-seq data (MOTIF scores see Supplementary Figure 2 and 3), do not necessarily reflect the *in vivo* activity of the variant enhancer (Figure 1D, E; red bars). This was particularly striking for the G3 site mutants, for which very different levels of activity were obtained although predicted affinity scores were undistinguishable. Thus, a-element *in vivo* enhancer activities are only partially explained by the *in silico* predicted affinities of the binding sites for their transcription factors. There was a high correlation between the *in silico* and the *in vitro* relative affinities of Ets1/2 and Gata4/5/6 for the mutated binding sites we tested, as determined by the quantitative multiple fluorescence relative affinity assay (QuMFRA) (Supplementary Figure 4, 5; 20). Thus, we conclude that altered *in vitro* affinity cannot provide a sufficient explanation for the observed effects of the point mutations on enhancer activity (Figure 1D and E).

Consistently, out of 14 genomic clusters of 2 ETS and 2 GATA sites with *in silico* predicted affinity scores for their cognate factor at least as good as the a-element, only two, N83 and N26, behaved as enhancers (Supplementary Table 1). The activity of these two elements was restricted to the early neural lineages. Conversely, out of 19 clusters tested by Khoueiry and colleagues (8), inactive cluster C39 has higher *in silico* scores for all its transcription factor binding sites than those of the a-element, while active C35 has 3 weaker scoring transcription factor binding sites. Thus, high *in silico* predicted octamer transcription factor binding sites affinity is not sufficient to explain neural enhancer activity of genomic clusters of ETS and GATA sites.

### Spacer sequences strongly affect enhancer activity

As neither the arrangement (8) nor the sequences of transcription factor recognition sites fully explain enhancer activity, we next tested whether the stretches of nucleotides located between transcription factor binding sites, subsequently called spacers, affect enhancer activity. We constructed a library of synthetic enhancers: each randomized variant shared with the a-element the six bases centered on the “GATA” and “GGAA” core sequences of the four GATA- and ETS-binding sites, respectively, as well as the orientation and spacing of these sites. All spacer sequences were, however, fully randomized, the four bases being equiprobable at each position. The *in vivo* enhancer activity of 34 randomized a-elements (Supplementary Figure 6A; Supplementary Table 2) was determined by electroporation as above.

While a large majority of naturally occurring genomic clusters of putative ETS and GATA sites are inactive, 25 out of 34 randomized a-element variants had an activity higher or equal to 10% of the wild-type activity, and were considered active (Figure 2). These enhancers displayed a wide range of activity levels with 11 of the variants being at least as active as the original a-element. Spacers are thus quantitative regulators of enhancer activity. The activity of these variants was mostly restricted to a6.5 and b6.5 lineages. In addition, most variants with higher activity than the WT enhancer, as well as three variants with lower activity, showed weak activity in other cell lineages, in which the a-element is weakly active, mainly neural plate and muscle cells (Figure 2 and Supplementary Figure 7). As expected, inhibition of the FGF-signalling pathway by treatment of electroporated embryos with the MEK inhibitor U0126 starting from the 16-cell stage, led to a loss of activity of the variant enhancers (Supplementary figure 8).

Taken together, these experiments establish a crucial role for spacer sequences in enhancer activity: they quantitatively modulate a-element enhancer activity levels, while qualitatively preserving spatial responsiveness of the element to the FGF pathway.

**Figure 2:**
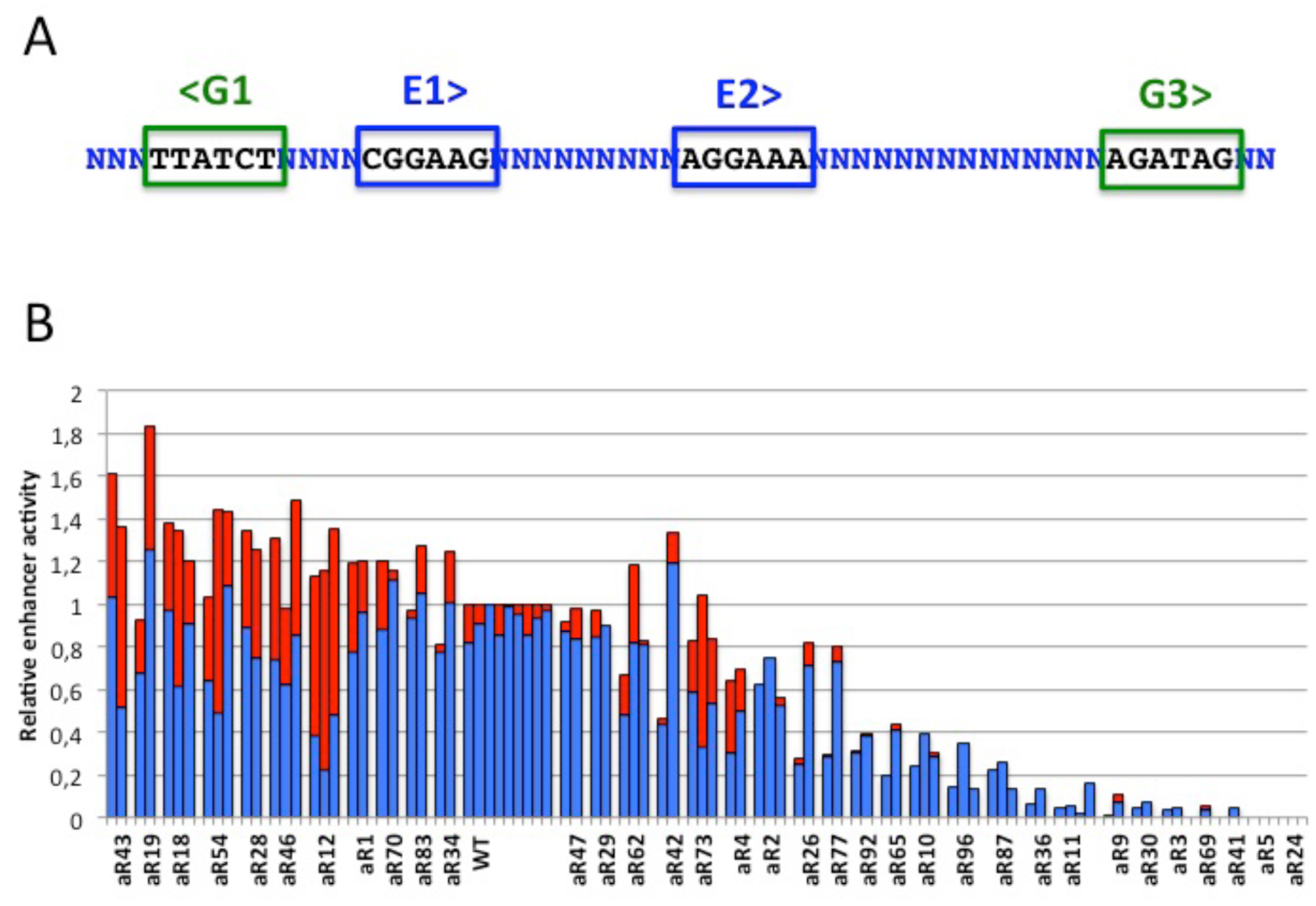
Randomizing the spacer sequences have major effects on the enhancer activity of the a-element of the Otx enhancer. A) The sequence of the a-element with randomized bases represented by N and the conserved GATA motifs boxed in green and ETS motifs in blue, the arrow pointing to their orientation. B) In vivo enhancer activity for 34 randomized variants normalized by the wild type enhancer activity in matching experiment. At least two independent experiments are shown for each construct. Blue bars correspond to exclusive expression in a6.5 and b6.5 lineages, red bars to additional expression in a6.7, B6.4 and other lineages.

### Randomized a-element variants with low *in vivo* activity have decreased *in vitro* affinity for Ets1/2 and Gata4/5/6 compared to high activity variants

To assess the influence of spacer sequences on the binding of Ets1/2 and Gata4/5/6 to the a-element we turned to *in vitro* binding experiments. We selected 9 randomized variants with equal or higher activity levels than the a-element and 9 variants with undetectable or very low activity (Figure 3A). Using the QuMFRA assay (21), we determined the relative *in vitro* binding affinities of the transcription factors for each of the complete enhancer variants (Supplementary Figure 4). Active variants showed significantly higher *in vitro* binding for the combination of Ets1/2 and Gata4/5/6 proteins than low activity variants (paired t-test, p=0.001) (Figure 3B). Both Ets1/2 and Gata4/5/6 proteins individually contributed to the binding preference for active enhancers, with a significantly higher contribution for Ets1/2 (paired t-test, p-value of 0.02327 for Ets1/2 compared to 0.05347 for Gata4/5/6; Supplementary Figure 9B).

**Figure 3:**
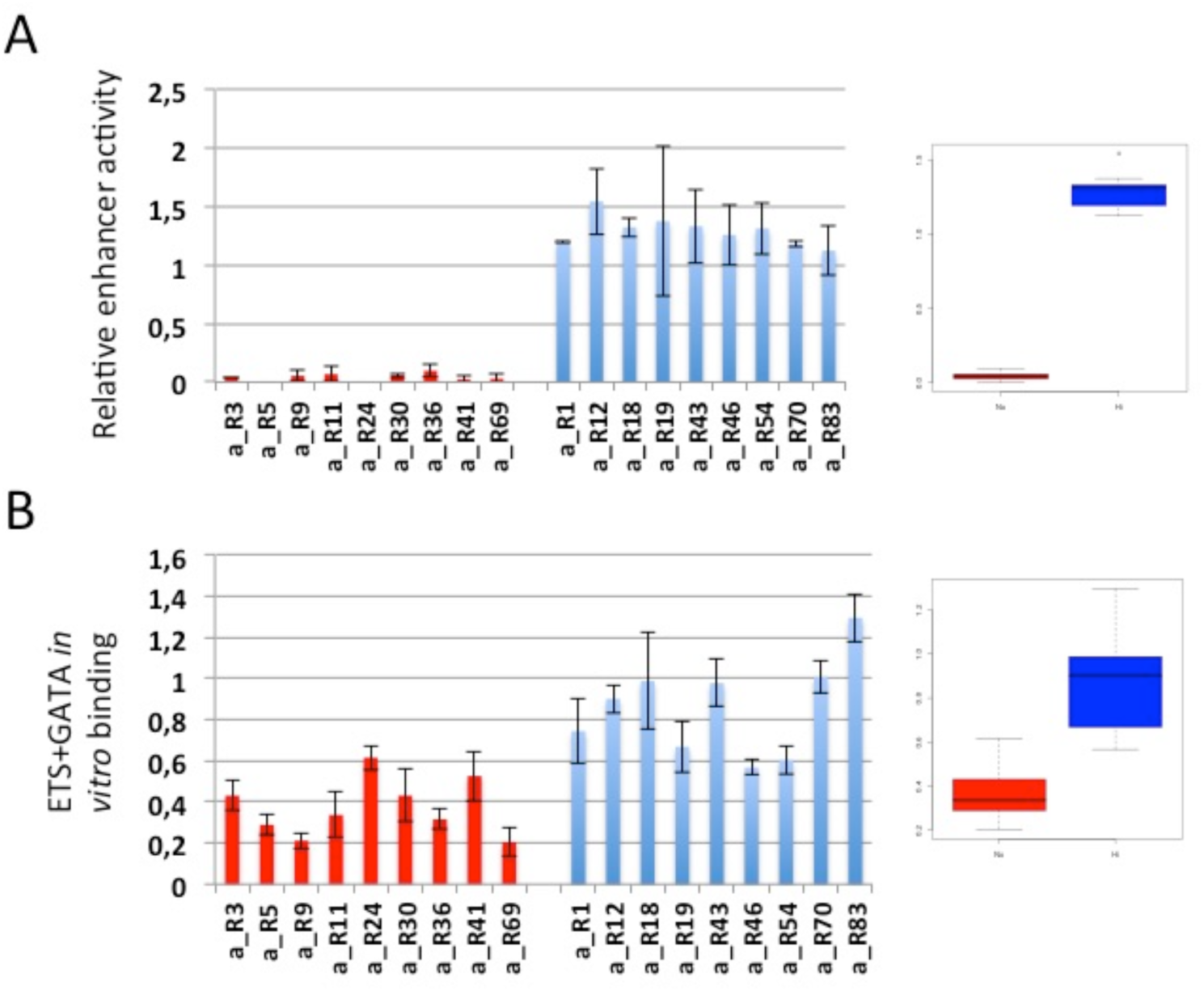
Active randomized enhancer variants have a higher *in vitro* affinity for Ets1/2 and Gata4/5/6. A) Relative *in vivo* activities of variant a-elements compared to the WT. Red: inactive variants. Blue: active variants. The right panel provides a populational description of activities. The two populations are different (paired t-test, p=2.825e-09). B) Relative *in vitro* binding of Ets1/2+Gata4/5/6 to the same variant a-elements as in A, compared to the WT. The right panel provides a populational description of binding. The two populations are different (paired t-test, p=0.001)

We conclude that spacer sequences affect the *in vitro* binding of Ets1/2 and Gata4/5/6 transcription factors to the enhancers. This could at least partly explain the wide range of enhancer activity levels obtained with the randomized variants.

### Flanking sequences modulate the affinity of Ets1/2 and Gata4/5/6 for their binding sites

We then analysed the individual sites in more detail in order to better understand the spacer effect on transcription factor affinity. First, we compared the binding of Ets1/2 and Gata4/5/6 to individual sites and to the complete enhancer of 9 variants (Supplementary figure 10 and Supplementary table 3). We note that binding to the individual sites mostly reflects binding to the complete enhancer, indicating that most information for binding of the transcription factors to the randomized variants is contained within the 30 bp sequence centered on their binding sites, with two exceptions: aR30 E1 site is a high affinity binding site although Ets1/2 is poorly binding to the complete enhancer, aR43 E1 and E2 sites are low affinity binding sites although Ets1/2 is strongly binding to the complete enhancer, possibly via a novel ETS site (E3) created in a randomized spacer sequence.

Farley and colleagues proposed that the activity of the enhancers is primarily determined by the two bases flanking each side of the “GGAA” or “GATA” core motif sequences (9). We note, however, that in our experiments active variant aR_30 and inactive variant aR_70 share very similar 8 bp-extended ETS and GATA sites, suggesting that additional bases of the spacers contribute to the spacer effect on affinity (Supplementary Figure 6A). To differentiate the effects of proximal bases immediately flanking the transcription factor binding sites to that of more distal effects, we focused on ETS sites, since the ETS sites displayed the most variable affinities.

We first tested a 10 bp binding site by keeping 10 bp of the randomized variants centered on the 6 bp core and completing the fragment with wild type flanking sequences to obtain a 30-mer (see oligonucleotide aRn-E1/E2s in Figure 4A). Ets1/2 demonstrated very different affinities for individual variants (Figure 4B). Importantly, sites with identical core octamers showed different affinities. For instance, the core octamer “TAGGAAAT” is present in both the high affinity aR9_E2 and the low affinity aR30_E2 sites, while the core “GCGGAAGG” is present in both the high affinity aR19_E1 and the low affinity aR5_E1 and aR43_E1 sites (Supplementary table 3). Thus, the observed affinities cannot be explained by the core octamer only, although the protein is not known to contact bases beyond this motif.

**Figure 4:**
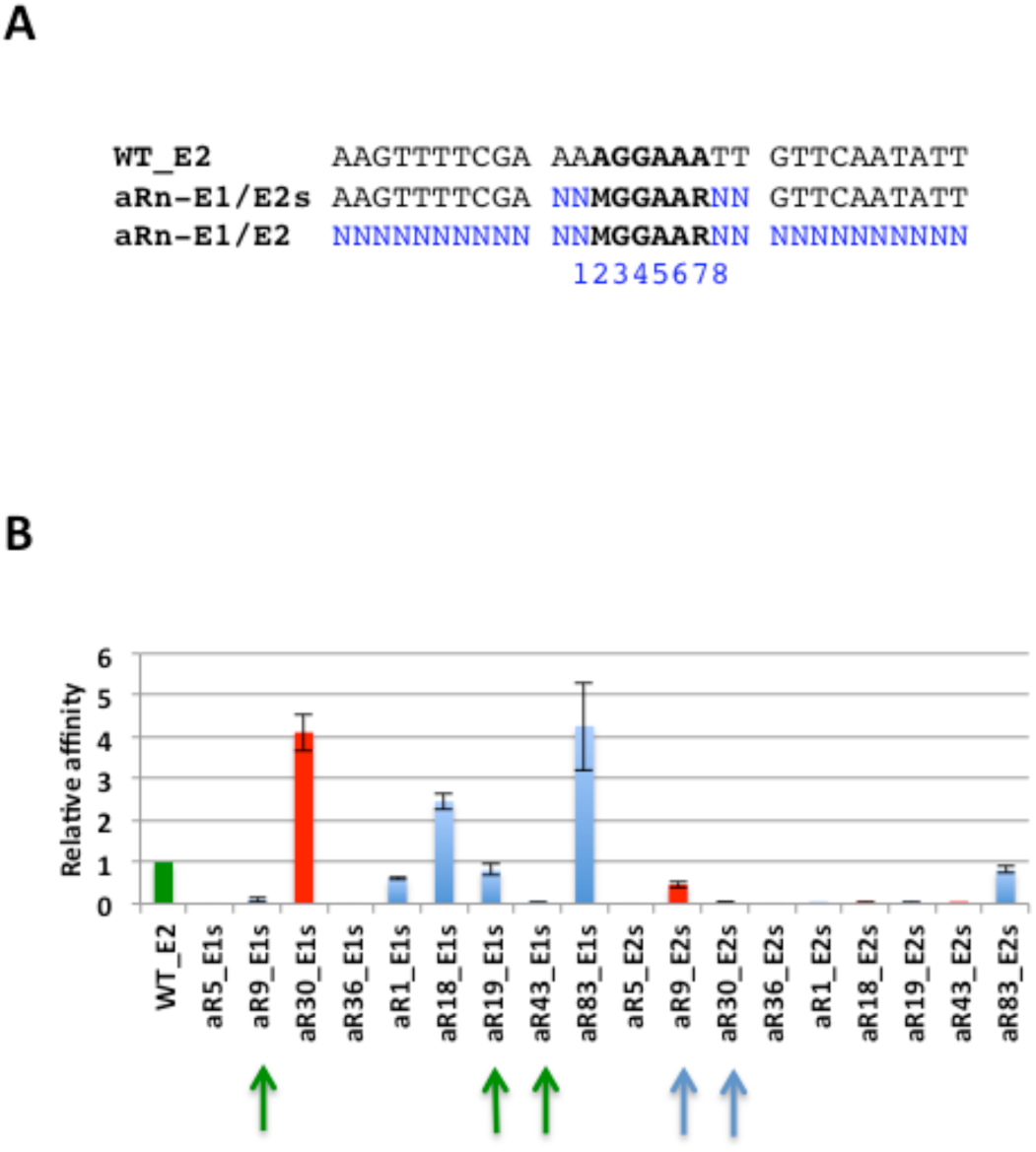
Sequences flanking the core octamer modulate the affinity for Ets1/2. A) Fragments used in the gel shift experiments. WT sequences are in black, randomized sequences are in blue. M is A or C, R is G or A, N is any base. B) Relative affinities of Ets1/2 for the transcription factor binding sites (aRn-E1/E2s) represented in (A) are with respect to in the 30 bp a-element fragment centered on Ets binding site E2 (WT-E2). Binding sites of active enhancers are in blue, binding sites of inactive enhancers are in red. Binding sites sharing the same octamer core are indicated with arrows of the same colour.

### Role of position −1 of ETS sites and shaping of the DNA helix

Consistently, mutation of the two base pairs flanking the common GCGGAAGG octamer of the aR5 and aR19 variants (Figure 5A) revealed that sites with a purine in position −1 display strong *in vitro* affinity for Ets1/2, while sites with a C in the same position have only weak or no detectable affinity. The identity of the base pair at position +9 was less important, although we noticed a decreasing affinity A>C>G>T.

**Figure 5:**
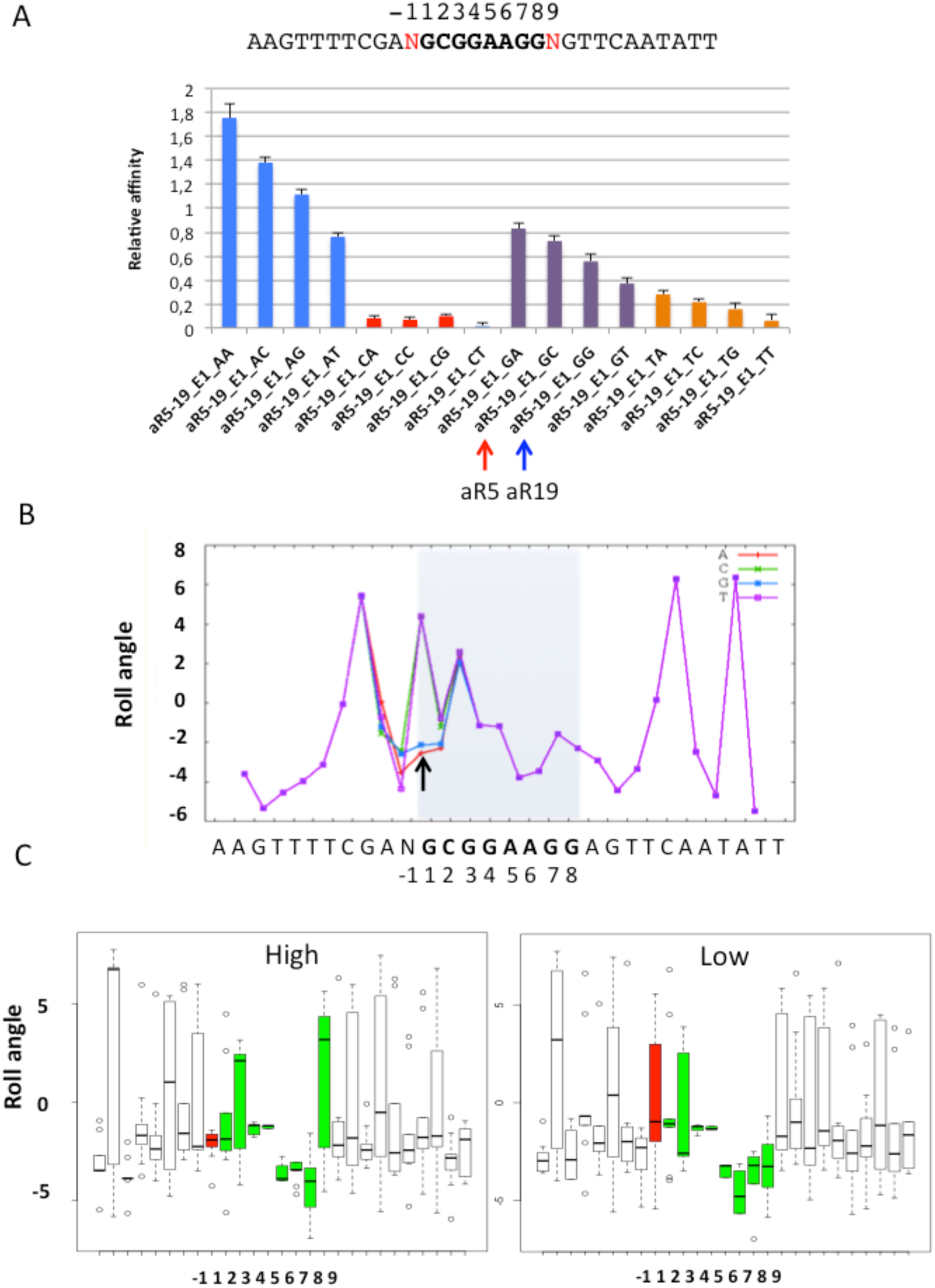
Essential role of position −1 in setting ETS affinity and in shaping the DNA helix. A) Effect on *in vitro* binding affinity to Ets1/2 of systematic mutagenesis of positions −1 and +9 (red) of aR5 and aR19 E1 binding site, which share a common octamer (bold). 10 bp flanking the E2 site of the WT a-element were added on either side of the test decamers. The red arrow points to the aR5 decamer, the blue one to the aR19 decamer. aR5-19-XY has an X in position −1 and a Y in position +9. Affinity is relative to WT-E2 as in Figure 4. B) Predicted roll angle with DNA shape for the oligomers with N = A, C, G or T in position −1 as indicated and A in position 9. ETS site is numbered as in figure 1C. Arrow indicates position between bp −1 and 1. C) Boxplot of predicted roll angles in degrees (Y-axis) for high affinity ETS sites left and Low affinity ETS sites right. Roll angles between consecutive base pairs between position 1 and 9 of ETS site in green. Roll angles between bases −1 and +1 are shown in red.

The crucial role of position −1 is further stressed by the fact that *in silico* affinity predictions based on Selex-seq data are improved by considering a 9-mer including the −1 position, instead of 8-mers (Supplementary figure 11). Addition of the +9 position does not improve the overall correlation (not shown).

Transcription factors recognize their target sequences by reading both the DNA sequence and the shape of the double helix (14). The failure of *in silico* prediction of transcription factor binding affinity to reliably account for enhancer activity when only the identity of the directly contacted base pairs are taken into consideration, suggests that the spacers may affect transcription factor binding distally through a change in the shape of the DNA helix over the TFBM nucleotides directly contacted by the Ets1/2 DNA binding domain (22). Interestingly, the presence of a purine at −1 leads to a negative roll between base pairs −1 and 1 as determined by DNA shape modelling, the roll representing the angle between two consecutive base pairs (Figure 5B) (21). All the high affinity tested sites had a negative roll, while the low affinity sites had a more variable distribution (Figure 5C). Thus, spacer sequences flanking the transcription factor binding site can modify the shape of the DNA helix at the transcription factor binding site, which may alter the affinity of Ets1/2 binding.

### More distal spacer sequences also affect the affinity of Ets1/2 binding

To assess whether the decamer bridging bp −1 and 9 is sufficient to explain the observed affinities or whether the more distally sequences can also contribute, we compared the *in vitro* affinities of the variant decamer ETS sites in the wild-type context (aRn-E1/E2s) to that of the complete variant 30-mer fragments (aRn-E1/E21 in Supplementary Figure 12). Indeed, further addition of spacer sequences at both sites of the decamer modifies the affinity of 5 of the 7 tested ETS binding sites (Supplementary Figure 12).

To analyze the relative roles of proximal and distal spacer sequences on transcription factor affinity, we replaced the E1 site decamer of the inactive aR5 variant by that of the active aR19 variant and *vice versa* and analyzed their *in vitro* interaction with Ets1/2 (Figure 6). The aR5 E1 core decamer conserved its low affinity in the aR19 context (aR5Ctx19) while the aR19 E1 core decamer in the aR5 context (aR19Ctx5) displayed a reduced affinity for Ets1/2 compared to its original context (Figure 6). Thus, while the identity of the two base pairs flanking the core octamer is important for the protein-DNA interaction *in vitro*, flanking sequences can further modulate a favourable combination.

**Figure 6:**
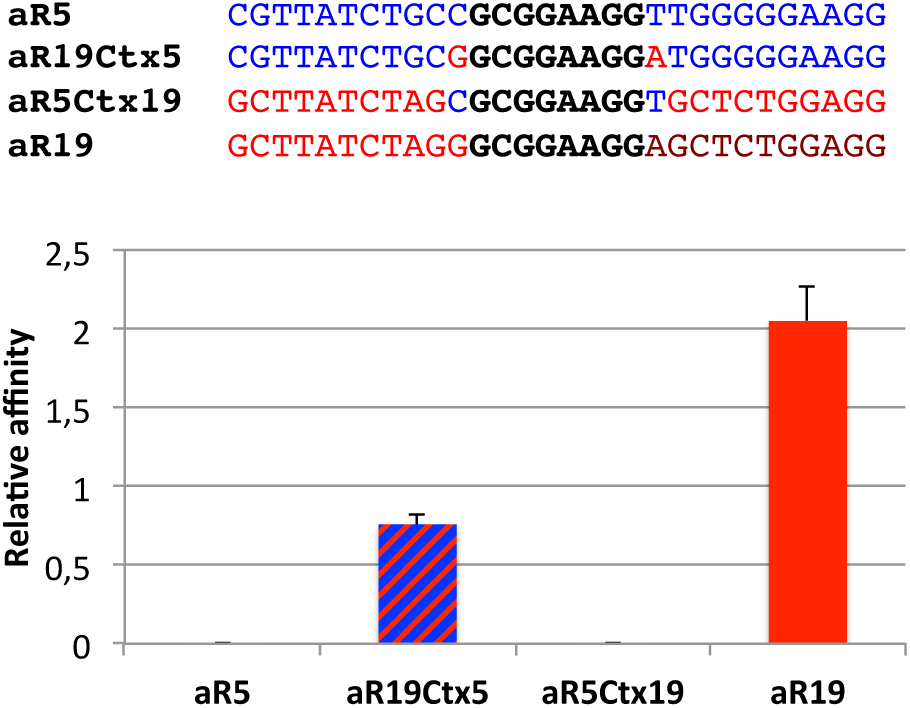
Role of sequences immediately flanking or more distantly located from the E1 core octamer. Comparison of *in vitro* Ets1/2 binding affinities to indicated decamers (Ctx=context). The sequences of the oligonucleotides used for the Gel Shifts are shown above the diagram. The invariant octamer core is in black, aR5 sequences in blue and aR19 sequences in red. Affinity is relative to WT-E2 as in Figure 4.

We conclude that although the two bases immediately flanking the Ets1/2 core octamer pairs can modify substantially the affinity of the factor, more distant flanking sequences also modulate this affinity.

### Activation of transcriptionally dormant clusters of ETS and GATA sites by randomization of spacer sequences

To test whether the major effect of spacer sequences on enhancer activity levels was a specific feature of the very compact a-element, we randomized the spacer sequences of a larger active genomic ETS and GATA cluster, N26 (Supplementary Table 2), whose sites spacing and orientation also differ from the a-element (Supplementary Figure 6B). Similar results (9/13 variants active in the neural lineages) were obtained with N26, suggesting that the contribution of spacer sequences to enhancer activity is not a specific property of the compact a-element (Figure 7A).

Most naturally occurring genomic clusters of ETS and GATA sites are transcriptionally inactive. To assess whether this inactivity could be due to inappropriate spacer sequences, we tested the effect of randomizing the spacers of two inactive genomic clusters. One, N61, has naturally occurring high scoring *in silico* recognition sequences for both factors. The second, C53 Opt, results from the optimization of the transcription factor binding sites of an inactive cluster C53. C53 Opt has itself no activity (8). Strikingly, spacer randomization conferred early neural enhancer activity to most variants of the two clusters (5/10 and 9/9 variants respectively) (Figure 7B, C; supplementary figure 6 C, D). We conclude that the inactivity of the tested genomic ETS and GATA clusters is due to inappropriate spacer sequences and that, very surprisingly, most randomized spacer sequences support enhancer function. This suggests that, although, by default, clusters of ETS and GATA binding motifs act as enhancers, most naturally occuring genomic clusters are kept inactive by inappropriate spacer sequences.

**Figure 7:**
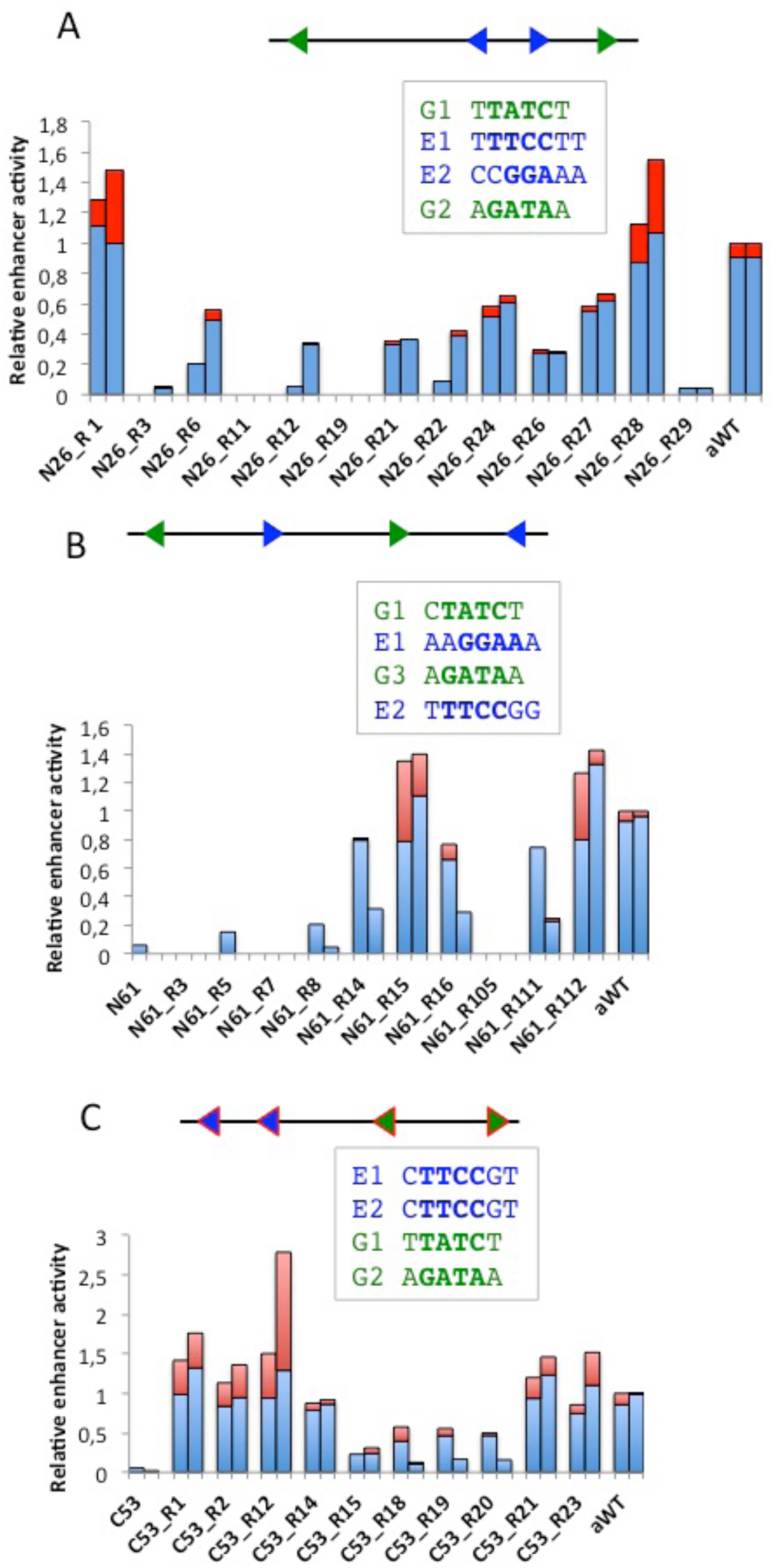
Randomization of spacer sequences activates inactive genomic clusters. Organization (top), and *in vivo* activity of three WT or randomized genomic clusters. A) Active genomic cluster N26. Variants R1, 6, 12, 21, 22, 24, 26, 27 and 28 are considered active. B) Inactive cluster N61. Variants 14, 15, 16, 111 and 112 are considered active. C) TFBM-optimized inactive cluster C53. All variants are active. For each set of elements, results of two independent electroporation experiments are shown. Blue and green arrowheads respectively represent the position and orientation of ETS and GATA sites, whose sequences appear below. Blue bars correspond to exclusive expression in a6.5 and b6.5 lineages, red bars to additional expression in a6.7, B6.4 and other lineages. The activity shown is relative to the activity of the a-element in the same experiment.

## DISCUSSION

In this study, we focused on the features able to confer activity to short clusters of ETS and GATA sites, and in particular to an early *Otx* neural enhancer in *Ciona intestinalis,* the a-element. Our previous work established that, in this element, the order and spacing of ETS and GATA sites was not a critical determinant of enhancer activity, yet that only a small minority of naturally occurring genomic clusters of such sites displayed enhancer activity (8). Thus, while dense clustering of transcription factor binding motifs has been proposed to explain enhancer activity (24), the fact that most genomic clusters of ETS- and GATA-binding motifs are transcriptionally inactive implies additional constraints. These could be provided by the chromosomal or chromatin environment (25) or by highly local sequence features independent of the binding motifs (25, 8). A repressive chromatin environment is unlikely to explain the lack of activity of most tested ETS and GATA clusters, since these short (<130 bp) elements were all tested outside of their normal genomic context and in the context of the same reporter vectors. Local features within the cluster sequences, and independent of TFBMs, were more likely (8).

These results prompted us to assay in the present work the activity of libraries of variants of clusters of ETS and GATA sites, in which individual bases of the TFBMs or the whole spacer sequences were systematically mutated. The main results presented here demonstrate that spacer sequences have a strong quantitative impact on enhancer activity, which in its magnitude is as strong as mutations in bases flanking the core nucleotides of TFBMs. Consistently, spacer activity can in part be attributed to modulations of *in vitro* binding of the transcription factors to their cognate TFBMs. This modulation involves bases located at a distance from the bases contacted by the transcription factors. These distal sequences may indirectly affect the shape of the DNA helix within the TFBMs. Finally, we found that in most cases, replacement of natural spacers by randomized ones suffices to confer enhancer activity to inactive genomic clusters of ETS and GATA sites, which may point to an unexpected mechanism to ensure that the majority of clusters born by chance in the genome are kept silent.

### A role for spacer sequences in the definition of enhancers in metazoan genomes

A recent report by Farley and colleagues analysed the activity of a large number of randomized variants of a slightly different version of the *Otx* a-element in *Ciona* (9) and concluded that the essential information for enhancer activity is included in octameric ETS and GATA sites. The authors did not detect a major role for spacer sequences, but this may reflect the different design of the two studies. Farley et al. only kept the four base pair core sequences for the two ETS and three GATA sites unchanged. The randomization thus affected both bases directly contacted by the transcription factors and directly flanking the core nucleotides, and the spacers as defined in our study. This strategy was therefore able to pick up both determinants of TFBM affinity to their transcription factors, which were the main interest of the authors, and spacer features. A closer analysis of their data indicates that sub-selections of their synthetic enhancers containing binding sites whose predicted *in silico* affinity for ETS and GATA is above a certain threshold, contain both active and inactive clusters with a wide range of activity levels. Thus, their data are consistent with a role for spacer sequences in setting enhancer activity levels.

In mammals, few studies involved the systematic large-scale analysis of enhancer variants. In most cases single point mutations were tested, and the resulting effects are expected to be less severe than the randomization approach described here and by Farley et al. (9). Evidence however also exists for a role of spacer sequences in enhancer activity. Kwasnieski et al. (27) reported the dramatic effects of base pair substitutions in the short 52 bp *rhodopsin* enhancer, some of which were located in spacer sequences. In larger enhancers, point mutations had a weaker effect, as reported by Melnikov et al. (28) and Patwardhan et al. (29) and these mutations mostly affected the transcription factor binding sites. It is thus unclear at present whether the impact of spacer sequences is restricted to short enhancers like the elements tested here or the mammalian rhodopsin enhancer, and/or whether they can also impact the activity of less compact elements.

### Spacer sequences can modulate the affinity of transcription factor binding sites by shaping the DNA helix

Our results suggest that spacer sequences affect enhancer activity by modulating the *in vitro* binding of transcription factors. Consistently, White et al. (25) showed that binding of the homeodomain protein Crx to clustered motifs was dependent on highly local sequence features such as high GC content. Parallel studies established that the affinity of transcription factors for specific DNA sites is controlled both by nucleotide sequence and DNA helix shape readouts, which are more permissive to variation in specific nucleotide sequence (30). In addition, a comprehensive computational analysis suggested that transcription factor binding is, in part, determined by nucleotides outside of the DNA sequence directly contacted (11, 12).

Our gel shift experiments provide experimental evidence suggesting that neighbouring base pairs can influence the specific interaction of Ets1/2 protein with the core motif of ETS binding sites by modifying the shape of the DNA helix. A similar observation was done for two yeast transcription factors by Levo et al. (22). Gata4/5/6 binding in contrast, seems to be less sensitive to local variations, which could reflect a greater flexibility of its zinc fingers compared to the alpha helices present in the helix-turn-helix motif of Ets1/2.

A finer dissection of the role of flanking base pairs of ETS binding motifs in setting the affinity revealed the importance of position −1 (Figure 5). Wei and co-workers mentioned the absence of a cytosine at this position in the group one Ets family, to which Ets1/2 belongs, and ascribed this to the unique presence in this group of a tyrosine in strand 4 of the DNA binding domain (19). Nevertheless, our Selex-seq data indicate that a cytosine is allowed in this position if it is followed by another cytosine. What could be critical is the presence of a negative roll between position −1 and 1, which is favoured not only by the presence of a purine at −1, but also by two consecutive cytosines. An intriguing possibility would be that a negative roll between bp −1 and 1, which opens the angle between the two base pairs, would facilitate base stacking interaction of the strand 4 tyrosine with the DNA helix. In addition, regions located distal to this extended binding site may also indirectly affect the DNA helix shape of the TFBS, as Ets1/2 *in vitro* affinity is further modulated by sequences outside of bases −1 to 7, a phenomenon particularly striking for variant aR30 (Supplementary Figure 10).

### Spacer sequences have other roles than modulating the affinity of the transcription factor binding sites

Altered *in vitro* binding of Ets1/2 and Gata4/5/6 to their binding sites in the randomized variants cannot provide an exclusive explanation for the variability in enhancer activity. Indeed, some inactive 55 bp variants are still binding Ets1/2 and Gata4/5/6 *in vitro* as efficiently as active ones (Figure 3B). Although this effect could in theory be explained by the creation in the randomized spacer sequences of binding sites for factors competing for Ets1/2 or Gata4/5/6 binding or for more general repressors of transcription, we did not find any correlation between potential novel sites and enhancer activity (not shown), suggesting additional mechanisms.

Enhancer sequences *in vivo* act within a complex chromatin environment, in which transcription factors and histones may compete for binding to DNA. Consistently, *in vivo* transcription factor binding assayed by chromatin immunoprecipitation only detects a minority of TFBMs able to bind their transcription factor *in vitro*. It is thus expected that spacers may not solely act by modulating the affinity of transcription factors to the naked DNA helix. Other processes potentially affected by spacer sequences could facilitate nucleosome exclusion (35, 8), DNA flexibility, which could help the formation of large TF complexes on the DNA, or allostery through the DNA helix, whereby fixation of one transcription factor facilitates binding of another transcription factor at a distance (33).

### Spacer sequences and the robustness of the transcriptional programme

One of the most unexpected results of our work is the facility with which a dormant ETS and GATA cluster can be awakened by spacer randomization, which seems in apparent contradiction with the small proportion of active naturally occurring clusters of high affinity ETS and GATA binding sites in the *Ciona* genome (8 and this study): although most synthetic spacers support enhancer function, the majority of natural spacers do not. This enrichment of "inactive spacers" in natural genomes may reflect the need to keep unwanted enhancer activity from appearing.

Creation, destruction or compensatory turnover of transcription factor binding sites is a frequent event (34), which could lead to the frequent appearance of clusters of transcription factor binding sites. In the *Drosophila* genome, positive selection contributes to transcription factor binding sites gain and loss, while purifying selection ensures their maintenance. Interestingly, the same trend was found in spacer sequences (35) and Parker et al. (36) provided evidence that the 3D shape of DNA is under selection in vertebrate regulatory regions. Similar selective forces could be at work in the compact genome of *C. intestinalis* (37), and explain that the majority of TFBM clusters feature spacers that ensure their inactivity.

Overall, the dependency of enhancer activity on a cross talk between transcription factor binding sites and spacer sequences could buffer against uncontrolled gene expression, thereby ensuring robustness of the developmental programme to sequence mutations.

## MATERIALS AND METHODS

### LacZ reporter assays

Mature *Ciona intestinalis* (type B) were provided by the Roscoff Marine Biological station and maintained in natural sea water at 16°C under constant illumination. Eggs were collected, fertilized and dechorionated as previously described (6).

Electroporation was performed as previously described (6) using the following parameters: 50μg DNA in 50μ1 H20 + 200μ1 D-Mannitol 0,96M; 50V-16ms pulse, using a Electro Square Porator machine (BTX T820; Harvard Apparatus). Embryos were grown in 0.1% gentamycin ASWH (Artificial Sea Water with Hepes) until their harvest at the 112-cell stage, where Xgal staining is found in every lineage that had expressed Otx so far, recapitulating the expression of the transgene at previous stages (6). Fixation and LacZ staining were performed as described in (6).

Where indicated, embryos were treated with a final concentration of 10μM U0126 from the early 16-cell stage (38). Control embryos were treated with the same amount of DMSO, added at the same time point (3μ1 DMSO in 15ml ASWH per plate). All experiments presented were at least repeated once.

At least 100 electroporated embryos where scored for each experiment by counting the % of embryos stained with LacZ in each territory, as this was shown to reflect enhancer activity (39). For each embryo, staining in a6.5 and/or b6.5 lineages, staining in other *Otx* expressing lineages (muscle, a6.7 cell lineage) and activities in territories not expressing *Otx* was retrieved. Enhancer variants driving detectable LacZ expression in less than 5% of stained embryos in all experiments were considered inactive. All values were normalized to the WT a-element activity electroporated in parallel in each experiment.

The level of activity in a given cell lineage is considered to be a function of the % of embryos in which X-gal staining is detected in this cell lineage.

### Gene IDs

*Ciona intestinalis Otx* : Gene model ID KH.C4.84 (Unique gene ID: Ciinte.g00006940)

*Ciona intestinalis Ets1/2*: Gene model ID KH.C10.113 (Unique gene ID: Ciinte.g00001309)

*Ciona intestinalis Gata4/5/6*: Gene model ID KH.L20.1 (Unique gene ID: Ciinte.g00012060)

### Construct design and molecular cloning

All these experiments were carried out using a modified version of the minimal wild-type element, in which a weak GATA binding site (G2; 8) is mutated without quantitative or qualitative impact on the activity of the element.

### Point mutations in ETS and GATA binding sites motifs

The family of GATA transcription factors preferentially binds the consensus “HGATAR” (H = A, C or T, R=G or A) (44; our SELEX data). Therefore, 12 variants of the a-element harbouring all variants of HGATAN at the third GATA position were designed to test both consensus (HGATAR) and non-consensus (HGATAY, Y = C or T) binding site motifs. The Ets family of transcription factors preferentially binds the consensus site “MMGGAWR” (M = A or C; W = A or T; R = G or A), with a higher affinity for “CCGGAWR” (45, 46), which is consistent with *Ciona* SELEX data (Nitta et al., in preparation). “T” at the seventh position was tested as negative control. 21 variants of the a-element were tested harbouring the different combinations MMGGAWD (D = A, G or T) at the second ETS site, with the exception of the MMGGATA combinations as they create an additional overlapping GATA site that could interfere with the ETS site activity and make the interpretation of the results not straightforward.

Oligonucleotides were synthesized containing part of Gateway attB1 and attB2 recombination sites in 5’ and 3’ respectively of the different elements we tested. These oligonucleotides were amplified by PCR using attB1F and attB2R primers, then inserted by successive BP and LR reactions in pDONOR221_P1-P2 and pDEST-L1 -RFA-L2-bpFOG-LacZ (43).

### Randomized variants

Only six bases per transcription factor binding sites, centered on “GGAA” and “GATA” for ETS and GATA respectively, were kept constant for the a-element variants. 7 bases were kept constant for ETS in randomized N26, N61 and C53 clusters, as it appeared that the 1^st^ base of the ETS site is important for *in vitro* binding.

Two nested PCRs were done to amplify the 5’ end of the insert containing an attB1 site, the sequence studied for its enhancer activity and the 5’ end of bpFOG using primers attB1F and P5 R first, then attB1 and P4 R. Three nested PCRs were performed to amplify the 3’ end of the insert containing the other half of bpFOG, the barcode and an attB2 site, using primers P1 F and attB2 R first, then P2 F and attB2 R then P3 F and attB2 R. Both fragments were assembled in a last PCR using attB1 F and attB2 R. They were then inserted in pDEST-L1- RFB-L2-LacZ by a one step BP-LR reaction (43)

### HT-SELEX

In SELEX assays, a tagged recombinant protein is incubated in solution with a degenerate mix of double-stranded oligonucleotides, comprising two constant ends of ~20 bases and a central portion of 12 to 24 random bases. The experiments analysed in this article were performed using oligonucleotides with 20 random bases and with bacterially-produced His-tagged transcription factor DNA-binding domains for the *Ciona intestinalis* ELK1/2/3 (nucleotides 314–818 of transcript model KH.C8.247.v2.A.SL3-1) and GATA4/5/6 (nucleotides 945–1319 of transcript model KH.L20.1.v1.R.ND1-1) proteins. The constant regions of the oligonucleotides contained a barcode, which was unique to each experiment and was used to multiplex oligonucleotide sequencing. The barcode included 6 bp 5’ of the randomized portion and 2–3 bp after the randomized region. Protein/DNA complexes were selected by chromatography on a Ni+-NTA sepharose (GE Healthcare) column recognizing the histidine tag. Bound oligonucleotides were then amplified by PCR using the constant ends of each oligonucleotide. The binding/chromatography/amplification steps were repeated for 3–7 cycles. After each cycle, the selected oligonucleotides were pooled and sequenced using Illumina Genome Analyzer IIx or HiSeq 2000 sequencer. Raw sequencing data were binned according to barcodes and used for further analyses. Unprocessed raw sequence data are available from the NCBI Short Reads Archive (SRA) (Accession XXXXXX). Before analysing the dataset, the constant region ends with the bar code were removed, leaving just the random portion of 20 bases. Duplicate oligonucleotides were removed from this set as they are most likely artefacts of the PCR amplification.

### In silico transcription factor binding site affinity prediction using MOTIF

We developed a software, called MOTIF, to estimate *in silico* the binding affinity of a transcription factor to a DNA sequence, based on SELEX-seq data (also called HT-SELEX), represented by the enrichment values of all 4096 6-mers present in the variable portion of the sequenced oligonucleotides bound to the transcription factor in the HT-selex procedure. The algorithm is as follows.

In a random set of oligonucleotides, k-mer frequencies will be distributed uniformly. HT-SELEX oligonucleotides are not random, as the method enriches for oligonucleotides bound to the transcription factor. The k-mer frequency distribution will thus become skewed, and the DNA-binding specificity of the transcription factor can be represented by an enrichment value for each of the k-mers considered. 6-mers were used here since 4,096 k-mers provides a sufficient number of k-mers without becoming sparse considering the depth of the sequencing. k-mer frequencies are determined by counting the occurrences of each k-mer in the set of unique oligonucleotides. To obtain an enrichment value, the observed count of each k-mer in the sequenced oligonucleotides, *obs,* are normalized using the expected count, *exp*, of each k-mer based on the number of sequenced oligonucleotides, *n*, with a variable oligonucleotide length of *d*, as shown in equation 1.

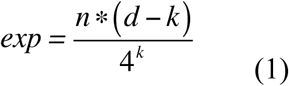

The enrichment score, *e*, was calculated as shown in equation 2.

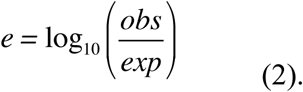

The synthesis method used to produce the original random pool of oligonucleotides is often biased, enriching certain k-mers over others. To correct for this bias, the enrichments are adjusted by the enrichment in the background set, shown in equation 3.

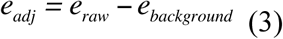

Many transcription factors recognize motifs longer than 6 bases. MOTIF thus associate to each base of the analysed DNA sequence a score predicting the binding of a transcription factor to the 8-mer starting at this base (Supplementary Figure 2). It corresponds to the sum of the 6-mer enrichments scores of the three 6-mers contained in each 8-mers.

### Selection of the14 ETS/GATA genomic clusters tested in vivo by electroporation

101 clusters containing at least 2 sites ETS and 2 sites GATA were identified in *Ciona intestinalis* genome, using SECOMOD, and a very relaxed consensus for the transcription factor binding sites sequences (as described in 8). We then looked for clusters of maximum 140 bp with at least 5 bp between two consecutive transcription factor binding sites. 55 conserved clusters were identified by (8). We tested the activity of an additional eight non-conserved and six conserved clusters in *C. savignyi*.

The 14 tested clusters contain 2 ETS and GATA sites with high MOTIF scores. Their sequences are listed in Supp. Table 1.

### 2-colour Fluorescent Electrophoretic Mobility Shift Assay

The DNA-binding domains of Ets1/2 (Ensembl ID: ENSCINT00000011848), i.e. aa 581–708, and GATA4/5/6 (Ensembl ID: ENSCINP00000009159), i.e. aa 291–415, were identified by homology to domains of orthologous human proteins used in the crystallographic 3D structure determination (Ets1, MMDB ID: 62790 and GATA1, MMDB ID: 106606). The corresponding DNA sequences were amplified by PCR from the cDNAs (44) and cloned in the expression vector pETG20A (EMBL Protein Expression and Purification Facility) by Gateway technology (Life Sciences). N-terminally poly-His-thioredoxin tagged recombinant proteins were produced in Rosetta-pLys-R strain and purified on Nickel Agarose columns as described in (45).

Enhancer DNA fragments were produced by PCR from the plasmids used for the LacZ reporter assays using Cy5 or Alexia 488-5’ labelled 19 nt primers (MWG Eurofins) flanking the enhancer sequences, i.e. TTGTACAAAAAAGCAGGCT for the forward and GGTACAATACACGAAGCTT for the reverse primer. DNA fragments containing unique transcription factor binding sites were synthesized directly (MWG Eurofins) and 5’-terminally labelled with Cy5 or Cy3 for the internal control on one strand.

The reaction conditions for the GS experiments were adapted from Hashimoto & Ware, (1995). Labelled DNA was incubated at 0.015 μM with recombinant Ets1/2 at 0.2 μM or Gata4/5/6 at 0.1 μM during 15 minutes at room temperature in 25mM Hepes pH7.9, 50mM KCl, 0.5 mM EDTA, 10% glycerol, 0.5mM di-thiothreitol and 100 μg/ml poly(dI-dC) and loaded on a 6% polyacrylamide gel in 0.5 % TAE, which was run at 10V/cm. The fluorescence was registered with an Amersham Imager 600 (General Electric) and quantified with the software provided by the supplier.

To have a better control over the experimental conditions we included an internal control: the randomized DNA fragments are fluorescently labelled with Cy5 and mixed with an equimolar amount of control DNA fragment labelled with Alexia 488 or Cy3. Relative affinities Y are quantified by reporting the fraction of shifted randomized DNA fragments to that of control fragment (21) (Supplementary Figure 4).

## ACKNOWLEDGEMENTS

We thank Dylan Da Cunha for help with the experiments and the Centre National pour la Recherche Scientifique, l’Université de Montpellier and l’Agence Nationale pour la Recherche (Grants Chor-Reg-Net ANR2005 NT05-2_42083; Chor-Evo-Net ANR2008 Blan 067 91; and TED ANR-13-BSV2-0011-01) for support. C. Cambillau, C. Dantec, P. Lemaire, J. Piette, U. Rothbächer, and R. Vincentelli were CNRS employees. M. Guéroult Bellone was supported by a doctoral contract from the University of Montpellier, K. R. Nitta and E. Jacox were supported by ANR grants to PL, a CNRS post-doctoral contract (KRN) and a Marie Curie Incoming International Fellowship (PIIF-GA-2010-272840, CisRegLogic, EJ).

## SUPPLEMENTARY DATA

**Supplementary Figure 1: DNA residues contacted by Gata3 and Ets1**

The logo is deduced from our Selex-seq data (for *C. intestinalis* Gata4/5/6 in (A) and Elk3 in (B). Contacts are deduced from the crystal structure-data of human Gata3-DNA complex (A; Chen et al. 2012) and these compiled by Wei et al. (2010) on mammalian Ets1-DNA complex (B). Conserved amino acids of *C. intestinalis* in contact with DNA are in bold; contacts with the base pairs are in black, contacts with water molecules in blue and with the sugar-phosphate backbone in green.

**Supplementary Figure 2: *In silico* k-mer binding affinity calculations with MOTIF**

Randomly synthesised oligonucleotides binding the *in vitro* produced DNA-binding domain of the transcription factors were enriched by the Selex-seq procedure. for each k-mer, its enrichment *e* was calculated as indicated in (B). The MOTIF score for each octamer, reflecting *in vitro* transcription factor binding affinity, is obtained by summing 3 consecutive 6-mer scores as shown in (A).

**Supplementary Figure 3: Moderate correlation between *in silico* binding predictions and *in vivo* activity**

A) a-element sequence describing ETS (blue) and GATA (green) site mutations tested *in vivo*. B) Comparison of *in vivo* enhancer activity and *in silico* predicted binding calculated for E2 octamer (MOTIF score). Each point corresponds to an ETS site variant individual *in vivo* experiment. C) Comparison of *in vivo* enhancer activity and *in silico* predicted binding calculated for G3. Each point corresponds to a GATA site variant individual *in vivo* experiment (triangles and circles respectively correspond to experiments 1 and 2 from Figure 1). Colours correspond to that of the last base and yellow circles correspond to the WT a-element.

**Supplementary Figure 4: Relative affinities are quantified by QuMFRA**

A) Gel shifts of the a-element with the DNA binding domains (DBD) of Gata4/5/6 (red), Ets1/2 (blue) separately (left) or combined (right). B-C) Quantification of the affinity of Ets1/2 and Gata4/5/6 for nine individual a-element mutants. Equimolar amounts of Att488 labelled WT a-element (B) and Cy5 labelled mutant or randomized variants (C) were incubated with recombinant Ets1/2 and Gata4/5/6 DBD and loaded on a 6% PAGE as explained in materials and methods. Panels B and C show the same gel imaged with the two wavelengths. The fluorescence of the shifted bands S and the total fluorescence T was quantified with a Amersham Imager 600. The relative affinity Y, shown on main figures, was calculated using the formula Y=(S_n_/T_n_)*(T_c_ /S_c_).

**Supplementary Figure 5: Comparison between *in silico* and *in vitro* relative binding affinities for tested mutants**

Relative *in silico* MOTIF scores (red bars) and *in vitro* binding affinities (purple bars) determined by gel shift assays for the indicated mutants. ETS is in (A) and GATA in (B).

**Supplementary Figure 6: DNA sequence alignment of randomized variants for the a-element, the active N26 cluster and the inactive N61 and C53 clusters**

DNA sequences were aligned using SeaView. ETS binding sites are represented by blue, and GATA binding sites by green arrowheads. a-element is in (A), N26 in (B), N61 in (C) and C53 in (D).

**Supplementary Figure 7: Some randomized variants have a broader activity pattern than the a-element**

Normalized *in vivo* enhancer activity is determined for WT and randomized variants of the a-element. The percentage of embryos where LacZ staining was only detected in a6.5 and b6.5 progeny is shown in blue, that where activity was detected in other cells appear in different colours corresponding to the cells drawn in dorsal and ventral views of a 112 cell-embryo. Activity in cells other than a6.5/b6.5 progeny was always associated with activity in a6.5 and/or b6.5 progeny.

**Supplementary Figure 8: Response of randomized variants to the FGF-signalling pathway**

Relative enhancer activities in control embryos or embryos treated with the MEK kinase inhibitor U0126 at the 16-cell stage. Enhancer activity relative to WT a-element is indicated in blue for a6.5/b6.5 only and in red for additional cells.

**Supplementary Figure 9: Comparison of *in silico* predicted and *in vitro* relative binding affinities of active versus inactive a-element variants**

A) Upper=Relative sum of the MOTIF score for the ETS binding sites compared to WT of the variants of figure 3 (paired t-test for the inactive versus active variants, p=0.06651). Lower=Relative sum of the MOTIF score for the GATA binding sites compared to WT of the variants of figure 3 (paired t-test for the inactive versus active variants, p=0.3789). B) Upper= Relative *in vitro* binding of Ets1/2 compared to the WT a-element of the same revertants as in (A). The two populations are different (paired t-test, p=0.01737). Lower=Relative *in vitro* binding of Gata4/5/6 compared to the WT a-element of the same revertants as in (A). The two populations are different (paired t-test, p=0.05347)

**Supplementary Figure 10: Affinity of Gata4/5/6 and Ets1/2 for isolated sites compared to the complete 55 bp a-element variants**

Relative *in vitro* transcription factor binding affinities are shown for the complete variants above the corresponding isolated GATA (A) or ETS (B) sites. Dark blue represents the relative contribution of the upper band corresponding to binding of two molecules in the gel shift experiments.

**Supplementary Figure 11: Nonamer scores are better predictors of Ets1/2 affinity than octamer scores**

*In silico* MOTIF Scores derived from the Selex-seq data were plotted against the *in vitro* relative Ets1/2 affinity determined by gel shift experiments with 30-mers centered on the GGAA motif. The scores were calculated for respectively octa- or nonamers as represented by boxes above the graphs.

**Supplementary Figure 12: Further addition of variant base pairs to the ETS decamer modulates the affinity for Ets1/2**

Comparison of relative *in vitro* Ets1/2 binding affinities of indicated E1 or E2 decamers in their original environment (l) or that of the E2 site of the a-element (s) as indicated above the diagram. Affinity is relative to WT_E2.

**Supplementary Table 1: Sequence and genome coordinates of genomic ETS and GATA clusters conserved (C) and non-conserved (N) in *Ciona savignyi* genome**

**Supplementary Table 2: Sequences of the a-element, N26, N62 and C53_Opt genomic clusters and their randomized variants**

**Supplementary Table 3: Sequences of the oligonucleotides used for the gel shift experiments**

